# Structural analysis of tilvestamab in complex with AXL

**DOI:** 10.1101/2024.12.04.626834

**Authors:** Eleni Christakou, Andrea J. Lopez, Gopinath Muruganandam, David Micklem, James B. Lorens, Petri Kursula

## Abstract

AXL is a receptor tyrosine kinase with a significant role in various biological processes and important medical implications, particularly in cancer. AXL transduces signals from the extracellular environment into the cytoplasm by binding to its ligand, growth arrest-specific protein 6 (GAS6). Activation of AXL leads to autophosphorylation of its intracellular domain and subsequent activation of downstream signaling pathways involved in cell proliferation, migration, differentiation, and survival. Tilvestamab (also known as BGB149) is a first-in-class, humanized, therapeutic anti-AXL function-blocking monoclonal antibody. We carried out a structural characterization of the AXL-tilvestamab complex, using both negative-stain and cryogenic transmission electron microscopy as well as synchrotron small-angle X-ray scattering. While AXL-Fc was highly elongated and formed large, heterogeneous complexes with the full antibody, homogeneous samples for structural studies could be made using the monomeric soluble AXL extracellular domain, the Fab fragment of tilvestamab, and an anti-Fab nanobody. Both SAXS and cryo-EM confirmed successful complex formation between the three proteins, and a low-resolution 3D model for the tilvestamab-AXL complex is presented. The data allow for sample optimization for high-resolution structural biology as well as designing mutations that could alter binding affinity and specificity.

## Introduction

AXL is a member of the TAM (TYRO3, AXL, MERTK) family of receptor tyrosine kinases (RTKs), which play critical roles in maintaining tissue homeostasis and regulation of the innate immune system^1,2^. In a disease setting, AXL signaling is associated with aggressive, drug-resistant, and immune-resistant tumor phenotypes^3,4^, fibrosis^5^ and may facilitate infection with a range of viruses^6^.

AXL is comprised of an extracellular ligand-binding domain with two immunoglobulin-like (Ig) and two fibronectin type III domains, a single transmembrane helix, and an intracellular tyrosine kinase domain^7^. AXL is activated by the growth arrest-specific protein 6 (GAS6)^8-10^, leading to activation of multiple downstream pathways depending on the cell type and context ^11^. The AXL Ig1 and Ig2 domains are important for GAS6 binding and thereby receptor activation. Earlier structural studies have provided insights into the AXL-GAS6 interactions necessary for activation^12^.

Due to its involvement in human disease, AXL has gained strong interest towards therapeutic and diagnostic applications, in addition to its importance to fundamental biology. In oncology, AXL overexpression is linked to poor prognosis, metastasis, and resistance to conventional treatments^13-16^. Consequently, AXL is considered a promising therapeutic target, with both small-molecule inhibitors^17-19^ and monoclonal antibodies ^20^ under development aimed at blocking its function. AXL is additionally implicated in fibrotic diseases and immune disorders, and it could function in internalization of viral particles, including SARS-CoV-2^21^, broadening the scope of its therapeutic and diagnostic potential.

Tilvestamab (BGB149) is a fully humanized monoclonal function-blocking anti-AXL therapeutic antibody, developed by BerGenBio. The tilvestamab-AXL interaction blocks the GAS6-mediated AXL activation, thereby inhibiting the receptor’s downstream signaling pathways involved in disease progression^20,22,23^. Details on the antibody-antigen complex at the molecular level would be of great benefit to further develop this tool and understand molecular mechanisms of disease and possible treatments.

We aimed at a structural characterisation of the complex between tilvestamab and AXL. Using both experimental negative-stain transmission electron microscopy (TEM), small-angle X-ray scattering (SAXS), and cryogenic electron microscopy (cryo-EM), as well as AlphaFold3 modelling^24^, we build a model for the AXL-tilvestamab complex that can further be used to understand binding determinants in the complex, the molecular mechanisms of functional inhibition, and the inherent flexibility of the AXL extracellular domain. Our findings provide a foundation for high-resolution experimental structure determination of the AXL-tilvestamab complex, with potential applications in treatment and diagnosis of AXL-related diseases.

## Materials and methods

### Preparation of protein samples

Human AXL-Fc containing the entire extracellular domain of human AXL was purchased from Evitria (Switzerland). Monomeric sAXL, comprising the soluble extracellular domain of AXL, was prepared by papain digestion, followed by protein A purification, as previously described^25^.

Tilvestamab (BGB149, BerGenBio, Norway), a humanized AXL monoclonal antibody ^20^ was obtained from BerGenBio. The Fab fragment of tilvestamab was prepared by papain proteolysis and purified using standard protocols. Briefly, 3 ml (60 mg) tilvestamab were dialysed against PBS overnight. Digestion buffer was prepared just before use (PBS + 20 mM cysteine HCl + 20 mM EDTA, pH 7). 1.5 ml of 50% immobilized papain slurry (Thermo Scientific, 20341) was pipetted to a 15 ml falcon tube using a cut pipette tip, and 12 ml of digestion buffer were added to the gel slurry. After centrifugation (300 rpm RT), washing was repeated, and washes were discarded both times. The gel was resuspended in 1.5 ml digestion buffer (1:1). 3 ml of dialysed tilvestamab were added, and the sample was shaken at 37 °C for 2.5 h. After digestion, the sample was centrifuged (3000 rpm, 4 °C) and supernatant was recovered. Another elution was done with 0.5 ml PBS. Supernatants were spun down and filtered through a 0.45-μm filter, before applying to the MabSelect SuRe column. The eluate from the column was analysed by SDS-PAGE.

The anti-NabFab nanobody^26^ cDNA was cloned into the pTH27 vector^27^ and expressed in *E. coli* BL21 Rosetta cells. Cells were grown in 0.8 L LB media supplemented with 100 μg/ml of ampicillin and 34 μg/ml of chloramphenicol at 37 °C until an OD of 0.5 was reached. Then, 1 mM IPTG was added, and the cells were grown for 20 h at 20 °C. Harvested pellets were resuspended with 20 mM Hepes, pH 7.5, 0.1 mg/ml of lysozyme, 0.5 mM TCEP, 10 mM Imidazole, and the cOmplete™, EDTA-free Protease Inhibitor Cocktail (Merck). The cell suspension was lysed by sonication and centrifuged at 20 000 g for 15 min at 4 °C. The supernatant was filtered through a 0.45 mm membrane (Sarstedt REF 831826) and loaded into a 3 ml of Ni-NTA agarose column, pre-equilibrated with equilibration buffer (20 mM Hepes pH 7.5, 250 mM NaCl, 0.5 mM TCEP) with 10 mM imidazole. The column was washed 4 CV with equilibration buffer with 10 mM imidazole and 8CV with 20 mM imidazole. The last wash of 9 CV with 20 mM Hepes pH 7.5, 100 mM NaCl, and 20 mM imidazole. The protein was eluted with 20 mM Hepes pH 7.5, 100 mM NaCl, and 350 mM Imidazole. The His-tag was cleaved with tobacco etch virus (TEV) protease during dialysis for 16 h in 20 mM Hepes 7.5 pH, 100 mM NaCl, and 1 mM DTT followed by reverse Ni-NTA chromatography. The eluted protein was gel filtered with Hiload 16/600 Superdex 75 column pre-equilibrated with 20 mM Hepes pH 7.5, 100 mM NaCl, and 0.5 mM TCEP. The quality of the purified protein was analyzed by sodium dodecyl sulfate polyacrylamide gel electrophoresis (SDS-PAGE).

### Negative stain electron microscopy

Prior to sample application, formvar/carbon grids with copper mesh (Electron Microscopy Sciences) were glow-discharged in an ELMO glow discharge unit for 30 s. Samples were diluted 1:10, 1:100 and 1:1000 for initial screening. For negative staining, a 2 µl drop of sample solution was absorbed to a freshly glow-discharged grid, washed with three drops of deionized water and stained with three drops of freshly prepared 1% uranyl formate. Micrographs were recorded at room temperature on a JEM-1400 electron microscope (JEOL) equipped with a LaB6 cathode and operated at 120 kV. Images were acquired with a 4096 × 4096 pixel CMOS TemCam-F416 camera (TVIPS) at a nominal magnification of 60000 and a corresponding pixel size of 2.29 Å under a defocus between 2.5 and 5.0 μm.

### Small-angle X-ray scattering in solution

Small-angle X-ray scattering (SAXS) data from AXL, tilvestamab, Fabs of tilvestamab and complexes of AXL and tilvestamab were collected on the SWING beamline of Soleil synchrotron (Saint Aubin, France) in HPLC mode ^28^. The data were collected using a wavelength (λ) of 1.033 Å and a sample-to-detector (EIGER X 4M) distance of 2.0 m, resulting in a momentum transfer (q) range of 0.004-0.5 Å^-1^ (q = 4π sin θλ^-1^; where 2θ is the scattering angle). This setup enabes samples to pass first through a SEC column and later into a capillary where they are exposed to X-rays and allows the collection of hundreds of scattering curves from the SEC peak that corresponds to the sample of interest. Prior to SEC-SAXS analysis, samples were centrifuged at 20798 g for 10 min at 4 °C. For each measurement, 50 µl of the sample was injected onto a BioSEC-3 300 or BioSEC-5 1000 column (Agilent Technologies) pre-equilibrated with PBS or 10 mM histidine, 150 mM NaCl, pH 6.

The flow rate was 0.3 ml/min. SAXS data were recorded with an exposure time of 990 ms per frame and a dead time of 10 ms between frames. Buffer data were collected at the beginning of the chromatogram. Data reduction, R_g_ evaluation over elution profiles, data averaging, and merging were performed using the beamline software Foxtrot (version 3.5.2). Subsequent final data processing and model building were carried out using the ATSAS package^29^, with CHROMIXS being used to obtain scattering curves for SEC-SAXS data^30^. Distance distributions were obtained using GNOM^31^, and *ab initio* models were built with DAMMIN ^32^ for dummy atom-based modelling, and MONSA for a comprehensive multi-phase modelling^29^. During the project, test data were also collected on the CoSAXS beamline at MAX-IV synchrotron (Lund, Sweden)^33^, but the data presented here are from SOLEIL, as they were all collected using the same beamline and settings.

### Cryo-EM sample preparation and data collection

0.25 mg of sAXL was mixed with Fab fragments from tilvestamab and the anti-NabFab nanobody in a 1:2:3 molar ratio. The sample was incubated on ice for 30 min and subjected to SEC using a Superdex 200 increase 10/300 column. Cryo-EM grids of the complex at 0.33 mg/ml were prepared by applying 3 μl to airglow-discharged 300 mesh copper C-flat R1.2/1.3 or Au-flat R 0.6/1 grids, which were then plunged into liquid ethane using a Vitrobot (FEI Mark IV).

19505 movies were collected at the iNANO centre (Aarhus University, Denmark) using a Titan Krios electron microscope (FEI) operated at 300 kV with a K3 Summit direct electron detector (Gatan) with GIF Energy Filter. Each movie was dose-fractionated to 53 frames with a dose rate of 1.2 e^-^/Å^2^/frame for a total dose of 59.6 to 60.5 e^-^/Å. The total exposure time was 1.4 s.

### Image Processing

The collected data were processed using cryoSPARC v4.7.034. After patch motion correction and patch CTF estimation and, manual curation 17,189 micrographs were selected. Initial particle picking was performed on denoised micrographs (3000 micrographs) using the Blob Picker job, followed by extraction (box size of 360 pixels and Fourier cropped by a factor of 2) and 2D classification. The resulting class averages were used for template-based particle picking with an increased box size (box size of 480 pixels and Fourier cropped by a factor of 2), followed by a second round of 2D classification.

Selected particles were subjected to *ab initio* reconstruction (three classes) and heterogeneous refinement. Particles were then re-extracted with a 480-pixel box size and refined using non-uniform refinement. Local refinement of the best 3D class improved density corresponding to a putative Ig domain binding region. The final map was sharpened with DeepEMhancer ^34^ (*tightTarget* setting).

### Model Building and Refinement

The atomic model of the tilvestamab Fab and the antiNabFab nanobody was constructed based on the cryo-EM structure 7pij^26^. The models of NabFab and the antiNabFab nanobody were fitted into the cryo-EM map using ChimeraX v.1.10^35^. The tilvestamab Fab sequence was then used to mutate the model in Coot v.9.8.93^36^. Namdinator^37^ was used to fit the initial atomic model to the density map and correct the model geometry. Subsequent refinement steps were performed iteratively in reciprocal space using Phenix v.1.21.1^38^. Finally, the AXL Ig1 domain was modeled using AlphaFold3 and fitted into the extra density using ChimeraX v.1.10.

### Data availability

All data are available online through Zenodo.org or dedicated databases (EMDB, PDB). Access codes and DOIs will be updated here upon publication.

## Results and discussion

### Imaging of tilvestamab and AXL using electron microscopy

Already from previous work, we knew that the dimeric AXL-Fc and monomeric sAXL are elongated and flexible^25^. For a first characterisation of AXL and tilvestamab samples for structural studies, we carried out negative stain EM imaging of the proteins alone and in complex. The antibody alone showed both monomeric and dimeric particles, while in complex with AXL-Fc, heterogeneous large particles were observed. While the latter confirmed complex formation, it also indicated high conformational flexibility and the formation of larger-order oligomers. This is not unexpected, due to the dimeric nature of both AXL-Fc and tilvestamab. Based on these data, we decided to move further with smaller fragments of both proteins, i.e. Fab fragments of tilvestamab and monomeric AXL extracellular domain, without the Fc dimerisation domain.

### sAXL-tilvestamab solution structures using SAXS

As the next step after characterizing the sAXL-tilvestamab Synchrotron SAXS data were collected in SEC mode for different combinations of AXL-Fc, sAXL, tilvestamab, Fab, and anti-NabFab nanobody. 3D models were built using DAMMIN for all components separately, and MONSA was run with simultaneous input of 5 scattering curves and 3 protein phases (sAXL, Fab, Nb). SAXS parameters are shown in **Table 1** and results in **Fig. 2-3**.

**Table 1.**
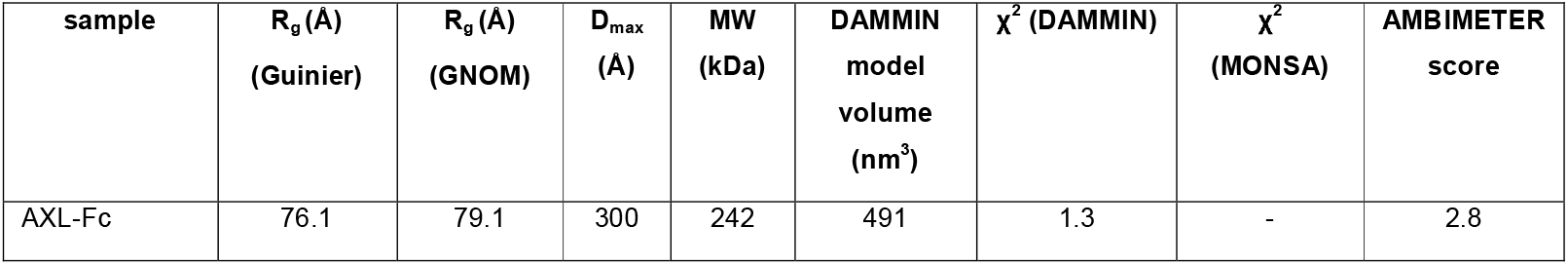
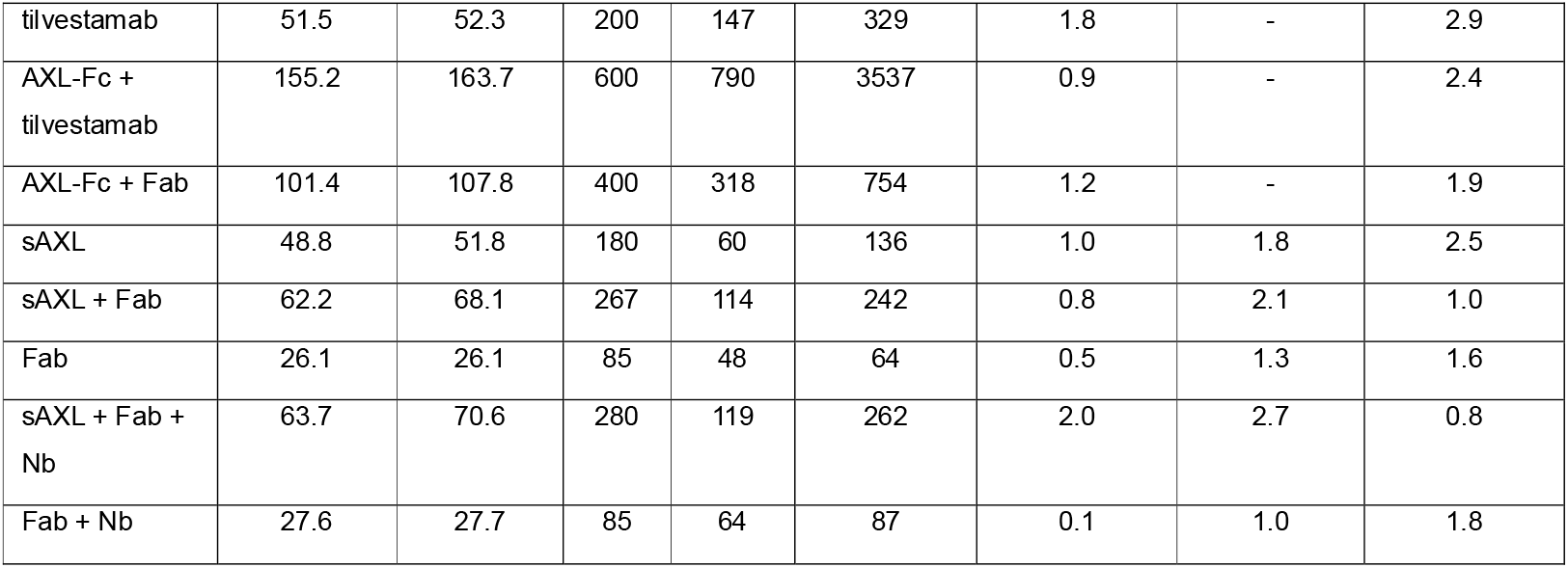
SAXS parameters. All data were collected using a SEC-SAXS setup. The MW corresponds to the Bayesian estimate given by PRIMUS. D_max_ comes from the distance distribution analysis in GNOM. *Ab initio* model volumes from DAMMIN are also shown, and they are affected by both MW, particle shape, and flexibility.

| sample | $R_g$ (Å)<br>(Guinier) | $R_g$ (Å)<br>(GNOM) | $D_{max}$<br>(Å) | MW<br>(kDa) | DAMMIN<br>model<br>volume<br>(nm<sup>3</sup>) | $\chi^2$ (DAMMIN) | $\chi^2$<br>(MONSA) | AMBIMETER<br>score |
| --- | --- | --- | --- | --- | --- | --- | --- | --- |
| AXL-Fc | 76.1 | 79.1 | 300 | 242 | 491 | 1.3 | - | 2.8 |
| tilvestamab | 51.5 | 52.3 | 200 | 147 | 329 | 1.8 | - | 2.9 |
| AXL-Fc +<br>tilvestamab | 155.2 | 163.7 | 600 | 790 | 3537 | 0.9 | - | 2.4 |
| AXL-Fc + Fab | 101.4 | 107.8 | 400 | 318 | 754 | 1.2 | - | 1.9 |
| sAXL | 48.8 | 51.8 | 180 | 60 | 136 | 1.0 | 1.8 | 2.5 |
| sAXL + Fab | 62.2 | 68.1 | 267 | 114 | 242 | 0.8 | 2.1 | 1.0 |
| Fab | 26.1 | 26.1 | 85 | 48 | 64 | 0.5 | 1.3 | 1.6 |
| sAXL + Fab +<br>Nb | 63.7 | 70.6 | 280 | 119 | 262 | 2.0 | 2.7 | 0.8 |
| Fab + Nb | 27.6 | 27.7 | 85 | 64 | 87 | 0.1 | 1.0 | 1.8 |

**Figure 1.**
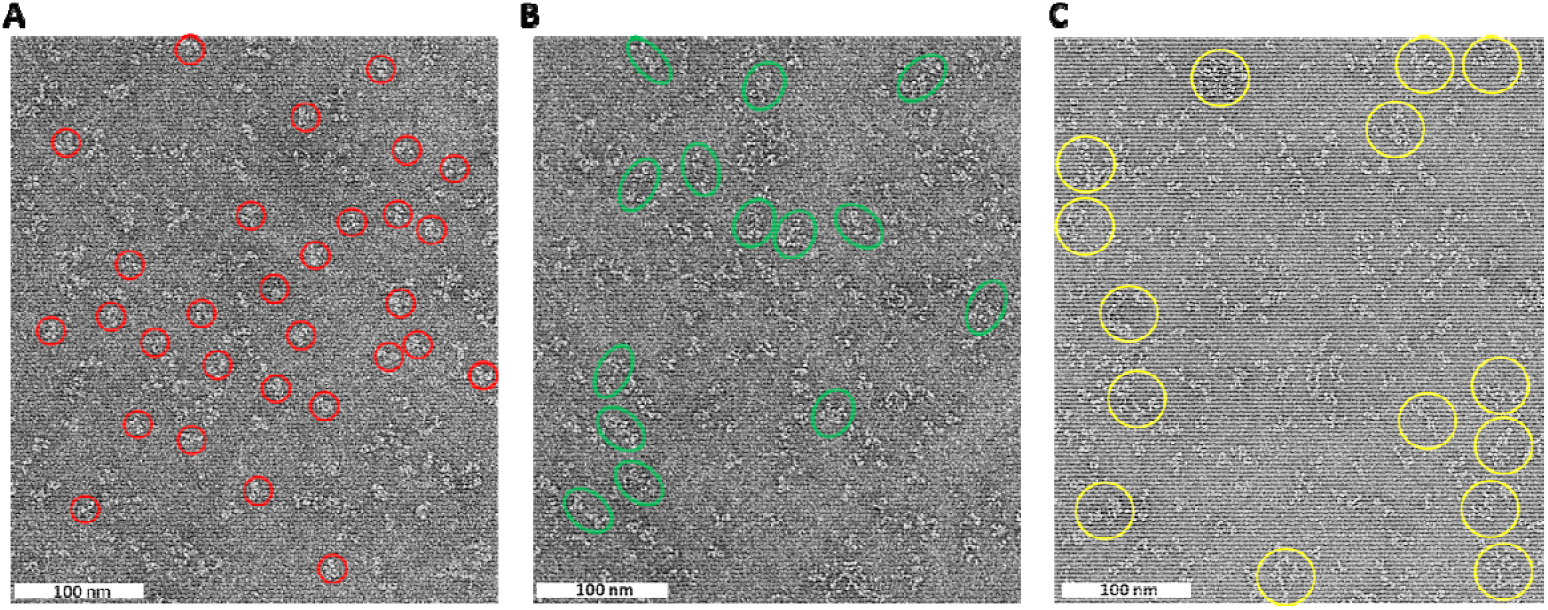
Negative-stain TEM analysis of AXL-Fc and tilvestamab. Representative micrographs of negative-stained molecules showing (**A)** predominantly monomeric tilvestamab particles marked with red circles, (**B)** an increased presence of dimeric tilvestamab particles shown as green ovals, (**C)** larger particles upon mixing tilvestamab with Axl-Fc circled in yellow. Scale bar: 100 nm.

**Figure 2.**
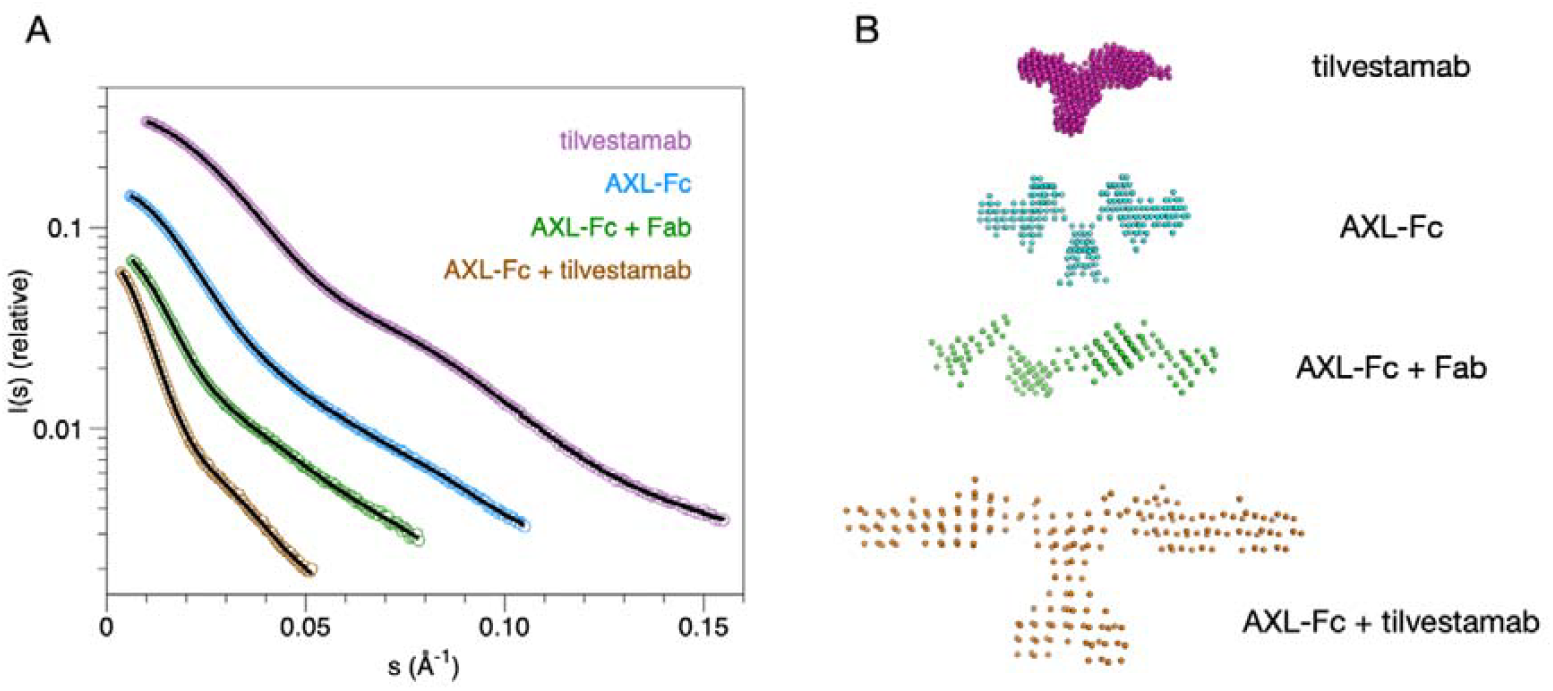
SAXS analysis of AXL-Fc and tilvestamab. (**A)** Scattering curves and DAMMIN model fits. (**B)** Dummy atom-based SAXS models of full-length AXL-Fc and tilvestamab. Guinier and Kratky plots as well as distance distributions are shown in Supplementary Figure 1.

**Figure 3.**
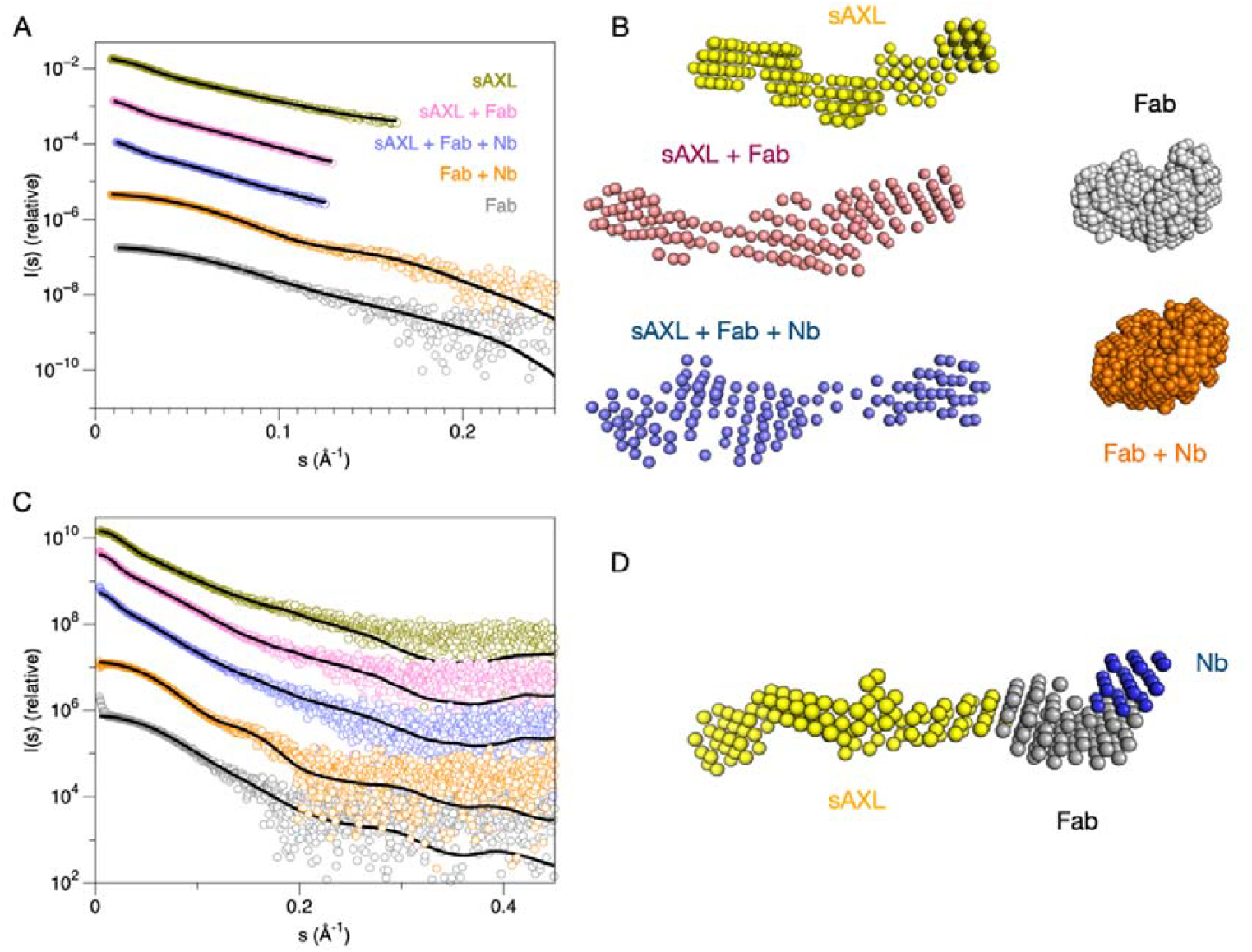
SAXS analysis for truncated variants. **(A)** Scattering curves and DAMMIN fits. (**B)** Dummy atom-based SAXS models for different combinations of sAXL, tilvestamab Fab, and anti-NabFab nanobody. (**C)** Fits to the same curves as in **A**, during multi-phase simultaneous fitting in MONSA. (**D)** A multi-phase model of the complex is shown, resulting from the simultaneous fitting of 5 different SAXS curves and 3 protein phases (sAXL, Fab, Nb). Guinier and Kratky plots as well as distance distributions and SEC-SAXS traces are shown in Supplementary Figure 1.

The AXL-Fc fusion was shown to be a very elongated, flexible dimer by SAXS, and mixing it with the full tilvestamab antibody resulted in large, heterogeneous complexes that could not be fully resolved even in SEC-SAXS (unpublished data, **Fig. 2**); these results agree with the negative stain EM results above. Therefore, the Fab fragment of tilvestamab was purified and used for additional SAXS experiments, to decrease sample heterogeneity.

AXL-Fc mixed with the Fab resulted in a heterotetrameric 2:2 complex, as expected (**Fig. 2**). This complex was even more elongated than the AXL-Fc dimer, indicating that the Fab bound close to the N terminus of AXL (the Fc fusion is C-terminal).

For further analyses, the Fc fragment was cleaved from the fusion, and the monomeric, soluble AXL extracellular domain (sAXL) was analysed alone and with tilvestamab Fab fragments. AMBIMETER^39^ scores calculated from each SAXS curve confirmed that the truncated sAXL as well as the Fab fragment were more homogeneous than the full-length proteins (**Table 1**).

sAXL alone was elongated, as previously seen^25^, having the four folded domains like beads on a string, similarly to the AlphaFold model. The Fab fragment increased the length and volume of the particle, and the MW corresponded to that of the expected 1:1 heterodimer (**Fig. 3**). Additionally, the anti-Fab nanobody was used to build a ternary complex, and the result indicated a further small increase in both R_g_, D_max_, and the molecular weight, showing that a 1:1:1 complex had been formed. The low-resolution structures from SAXS experiments therefore indicated an elongated, flexible structure for the AXL extracellular domain and confirmed the binding of the tilvestamab antibody to the first Ig domain of AXL. In an attempt to obtain higher-resolution data, we therefore moved into cryo-EM experiments with the ternary complex.

### Cryo-EM study for the AXL-tilvestamab Fab complex

Initial cryo-EM studies on dimeric AXL-Fc complexed with the tilvestamab Fab showed high degrees of flexibility of a very elongated, heterogeneous complex (unpublished data), in line with the SAXS data above (**Table 1, Fig. 2**). Therefore, based on the promising SAXS data (**Fig. 3**), monomeric sAXL was complexed with the tilvestamab Fab bound to anti-Fab nanobody for cryo-EM.

To obtain higher-resolution insights into the AXL–tilvestamab complex, particularly the epitope–paratope interactions, we prepared samples for cryo-EM analysis using monomeric sAXL, the tilvestamab Fab, and an anti-Fab nanobody (**Fig. 4, Table 2**). Based on SAXS data for the individual components and their complexes (**Fig. 3**), we anticipated structural flexibility of sAXL within the ternary complex. Cryo-EM reconstruction resulted in a map of the tilvestamab Fab bound to the anti-Fab nanobody at a global resolution of 3.66 Å (GS-FSC 0.143, soft mask), which improved to 3.09 Å with a tighter auto-generated mask. An additional density adjacent to the variable domains of the tilvestamab Fab was observed, which we attributed to the first Ig1 domain of AXL (**Figs. 5–6**).

**Table 2.** Data collection and refinement statistics.

| **Data collection** |  |
| --- | --- |
| Magnification | 130,000 x |
| Defocus range (mm) -Voltage (kV) | -300 |
| Microscope | Titan Krios |
| Detector | Falcon 3 |
| No. of frames | 53 |
| Pixel size (Å/pixel) | 0.647 |
| Electron dose (e<sup>-</sup>/Å<sup>2</sup>) | 59.6 |
| No. of micrographs | 19505 |
| **Reconstruction (CryoSPARC)** |  |
| No. of particles | 508,414 |
| Box size (px) | 480 |
| Average resolution (Å) (FSC=0.143) | 3.09 |
| Model resolution (Å) (FSC=0.5) | 3.03 |
| Map sharpening B-factor (Å<sup>2</sup>) | -90 |
| **Model building (Phenix 1.21.1_5286)** |  |
| No. of chains | 3 |
| No. of atoms | 4168 |
| No. of amino acid residues | 544 |
| No. of ligand atoms | 0 |
| Average B-factor (Å<sup>2</sup>) | 66.25 |
| R.m.s.d. bond lengths (Å) | 0.002 |
| R.m.s.d. bond angles (°) | 0.592 |
| CC volume | 0.75 |
| CC masked | 0.75 |
| Molprobity score | 1.35 |
| Clash score | 6.35 |
| Ramachandran plot favored/allowed/outliers (%) | 98.13/1.87/0 |
| Rama-Z (whole/helix/sheet/loop) | 0.38/1.33/0.33/0.41 |
| **Deposition codes** |  |
| PDB | - |
| EMDB | - |

**Figure 4.**
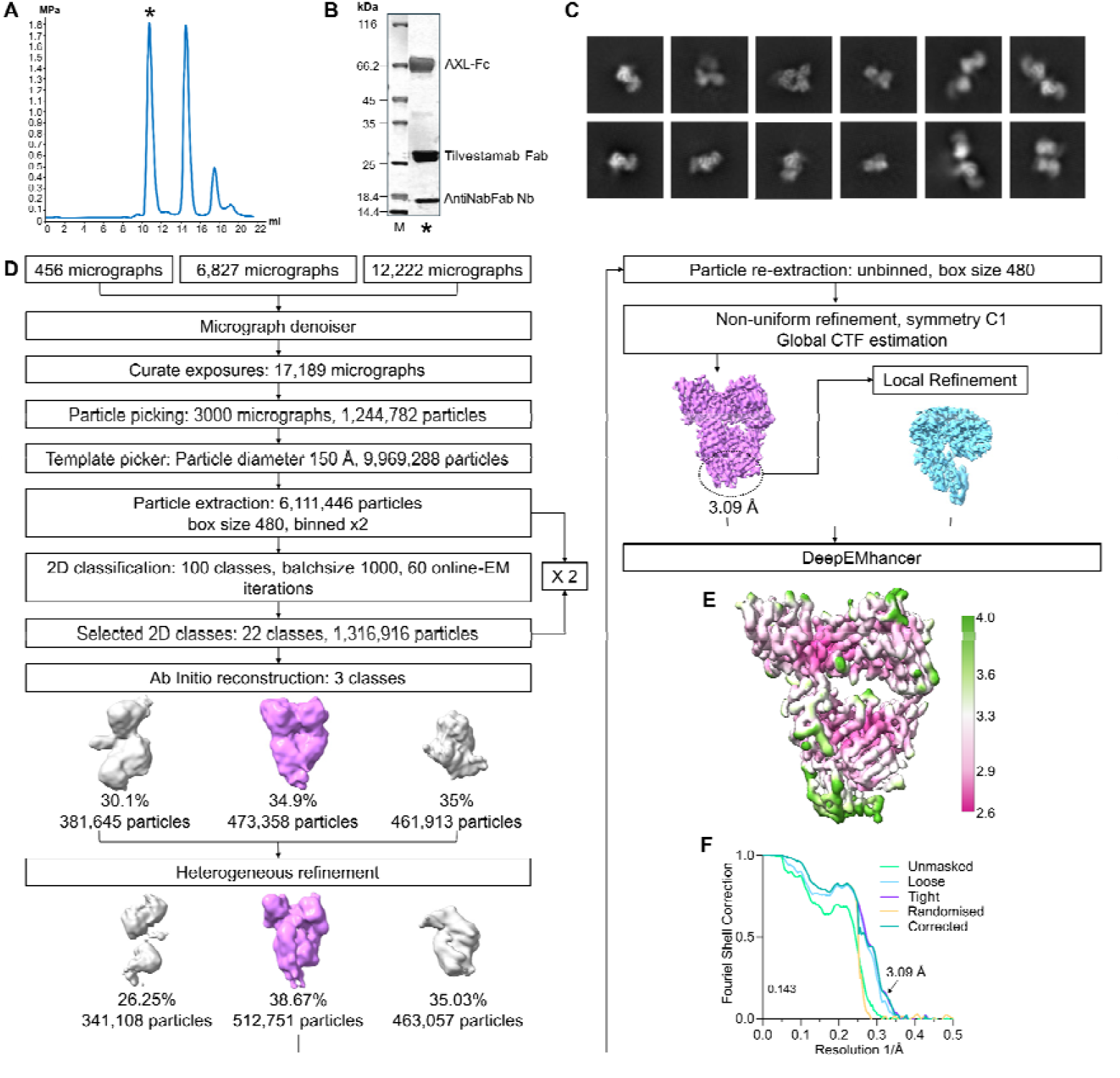
Cryo-EM sample preparation and processing. **A**. Size exclusion profiles of Fab, Axl-Fc, and the antiNabFab Nanobody (Nb). **B**. SDS-PAGE of the peak complex fraction (*) was used for sample preparation for cryo-EM. **C**. 2D classes. **D**. Schematic representation of the processing workflow. **E**. Final deepEMhancer sharpened map, colored according to local resolution, is displayed at 0.0817 σ. h. **F**. Fourier-shell correlation (0.143 criteria) of the final map.

**Fig 5.**
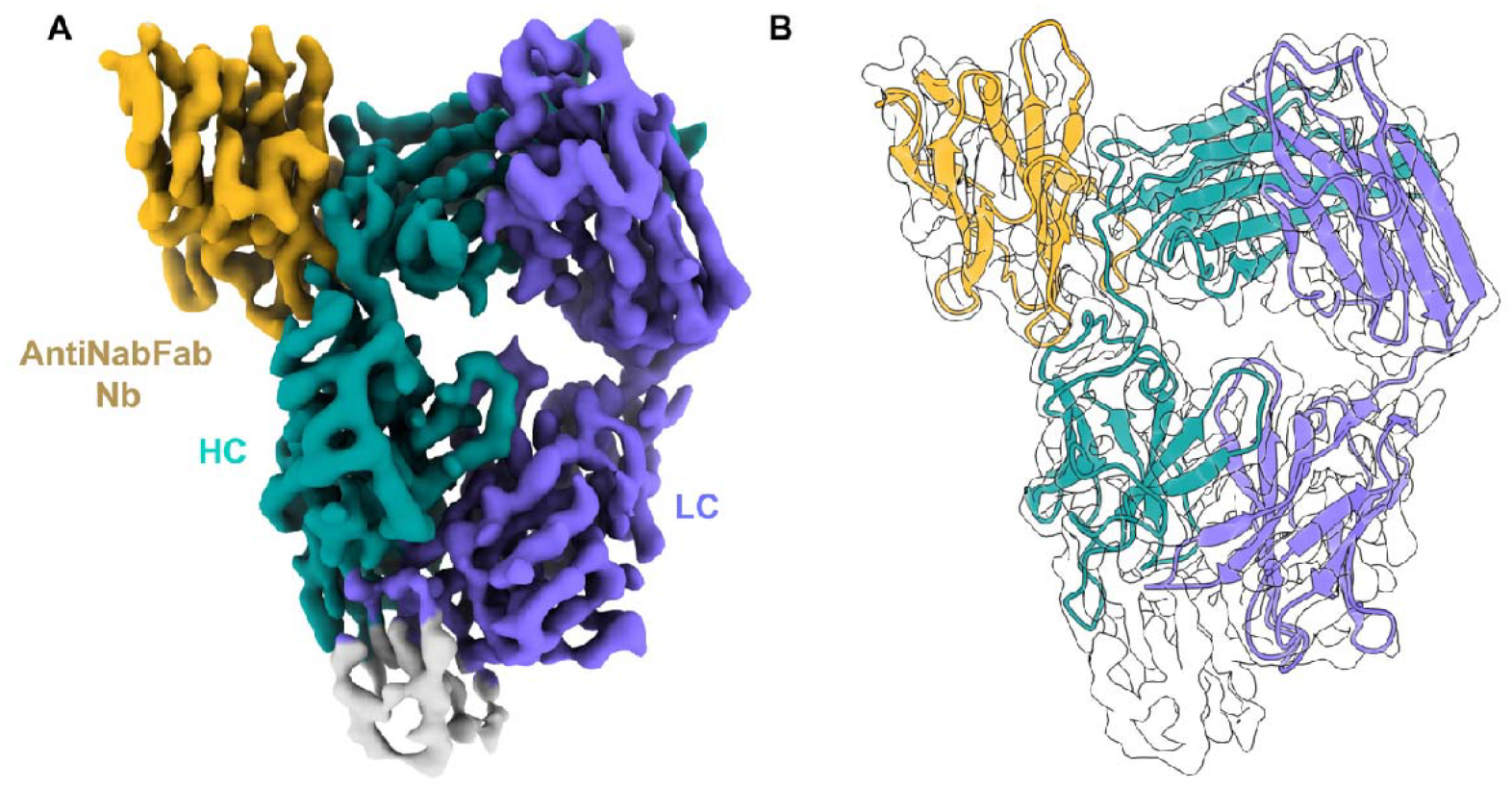
Reconstruction of Fab-Antinabfab complex. **A**. Final refined 3D map 3.09 Å resolution. Density for HC is shown in turquoise, purple for HL, and dark gold for the antiNabFab nanobody. **B**. Refined atomic model using Phenix fitted into the map.

**Figure 6.**
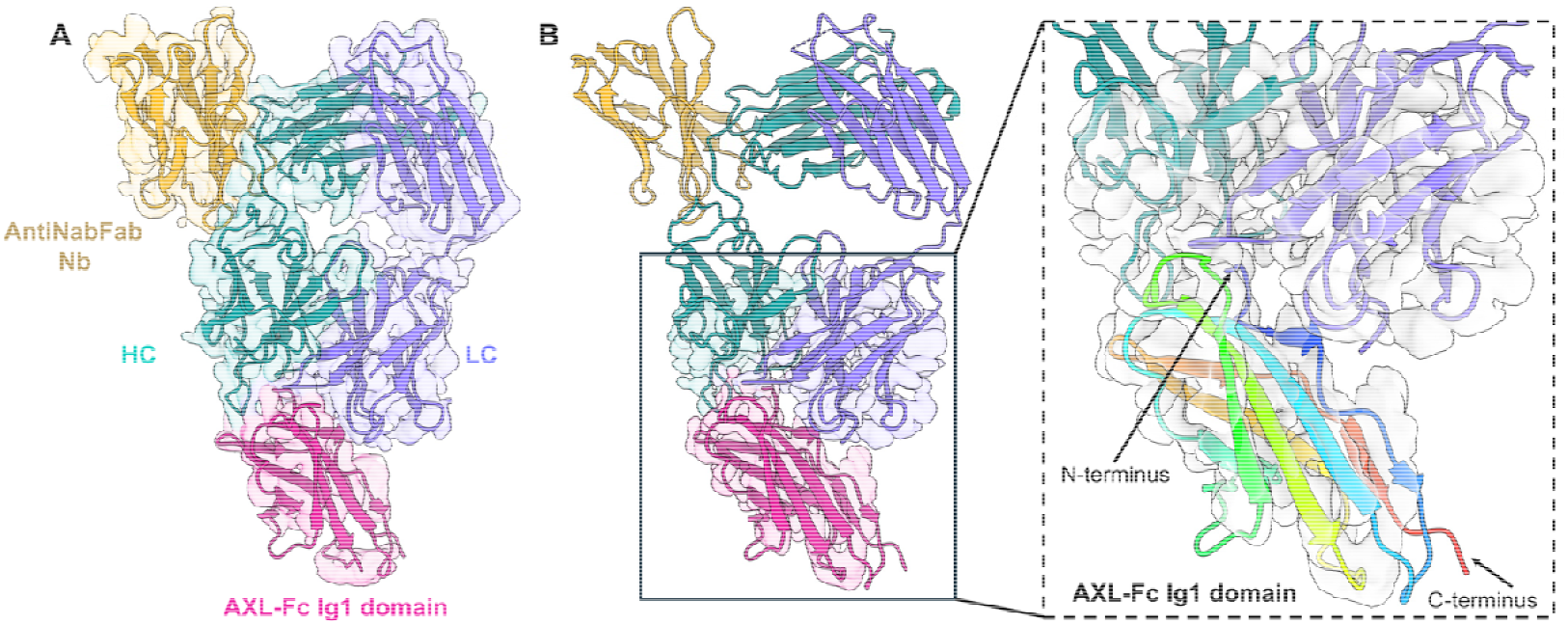
Fitting of the AXL Ig1 domain after local map refinement. **A**. Final refined 3D map. **B**. The local refinement clearly establishes the orientation of the antigen AXL Ig1 domain with respect to the antibody.

Comparison of the cryoEM map to the AlphaFold3 model (**Fig. 7**) shows that while the Fab-Nb complex is well in place and the binding interfaces on both AXL Ig1 and tilvestamab are correctly identified by the prediction, the orientation of the AXL Ig1 domain is flipped by approximately 180° between the experimental map and the model. This indicates that, as already known for antigen-antibody complexes^24,40^, AI-based methods may sometimes fare poorly in predictions of antibody-antigen interactions, and experimental methods still are crucial for deciphering the exact binding mode between the paratope and epitope.

**Figure 7.**
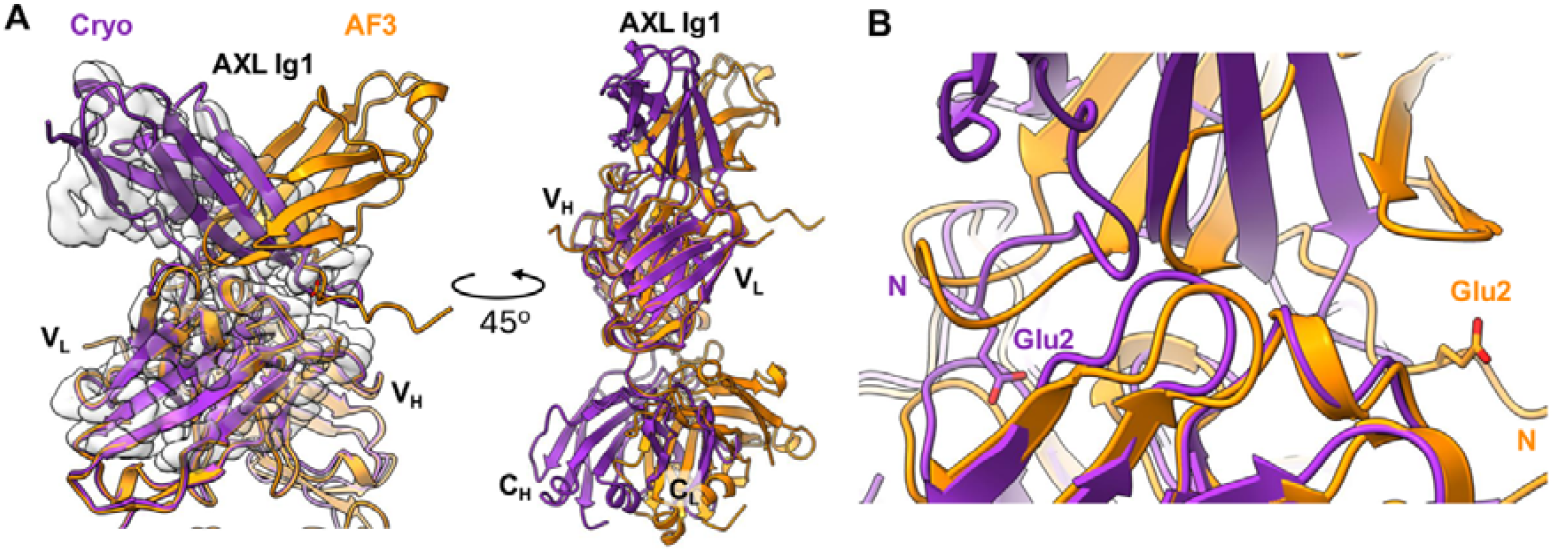
Comparison of Cryo-EM structure and AlphaFold3 prediction. **A**. Superimposition of the cryo-EM structure (purple) and the AlphaFold3 (AF3) predicted model (orange), fitted into the locally refined cryo-EM map (as in Fig. 7B). Alignment was performed using the V_L_ domain of the cryo-EM model as reference. In the AF3 prediction, the Ig1 domain of AXL exhibits a 180° rotation relative to the experimental structure, indicating correct identification of the binding surface but an incorrect domain orientation. **B**. In the AF3 model, the C_L_ and C_H_ domains are tilted approximately 30° towards the AXL Ig1 domain compared to the cryo-EM structure, suggesting a deviation in interdomain orientation. **C**. Both experimental and predicted models show Glu2 of AXL Ig1 directly involved in the interaction, implying that AF3 accurately identified the interaction interface despite misorienting the Ig1 domain.

As tilvestamab was characterised to block signalling along the GAS6-AXL axis, we compared the cryo-EM model to the crystal structure of AXL Ig1-2 with GAS6^12^. It becomes evident that the binding sites for GAS6 and tilvestamab are next to each other, with a small overlap (**Fig. 8**). This can explain, why tilvestamab does not fully block GAS6 binding^22^, while it fully inhibits downstream signalling^20,22,23^. The effect is likely a combination of weakening of GAS6-AXL interactions, conformational changes, and prevention of the formation of functional GAS6-AXL receptor complexes with correct AXL dimerisation in the context of the full-length proteins. Antibody-mediated prevention of dimerisation was suggested before for pembrolizumab and its target PD-1 receptor^41^. It is important to remember that this analysis is based on cryo-EM and crystal structures of truncated proteins, and conformational aspects of full-length proteins are not captured.

**Figure 8.**
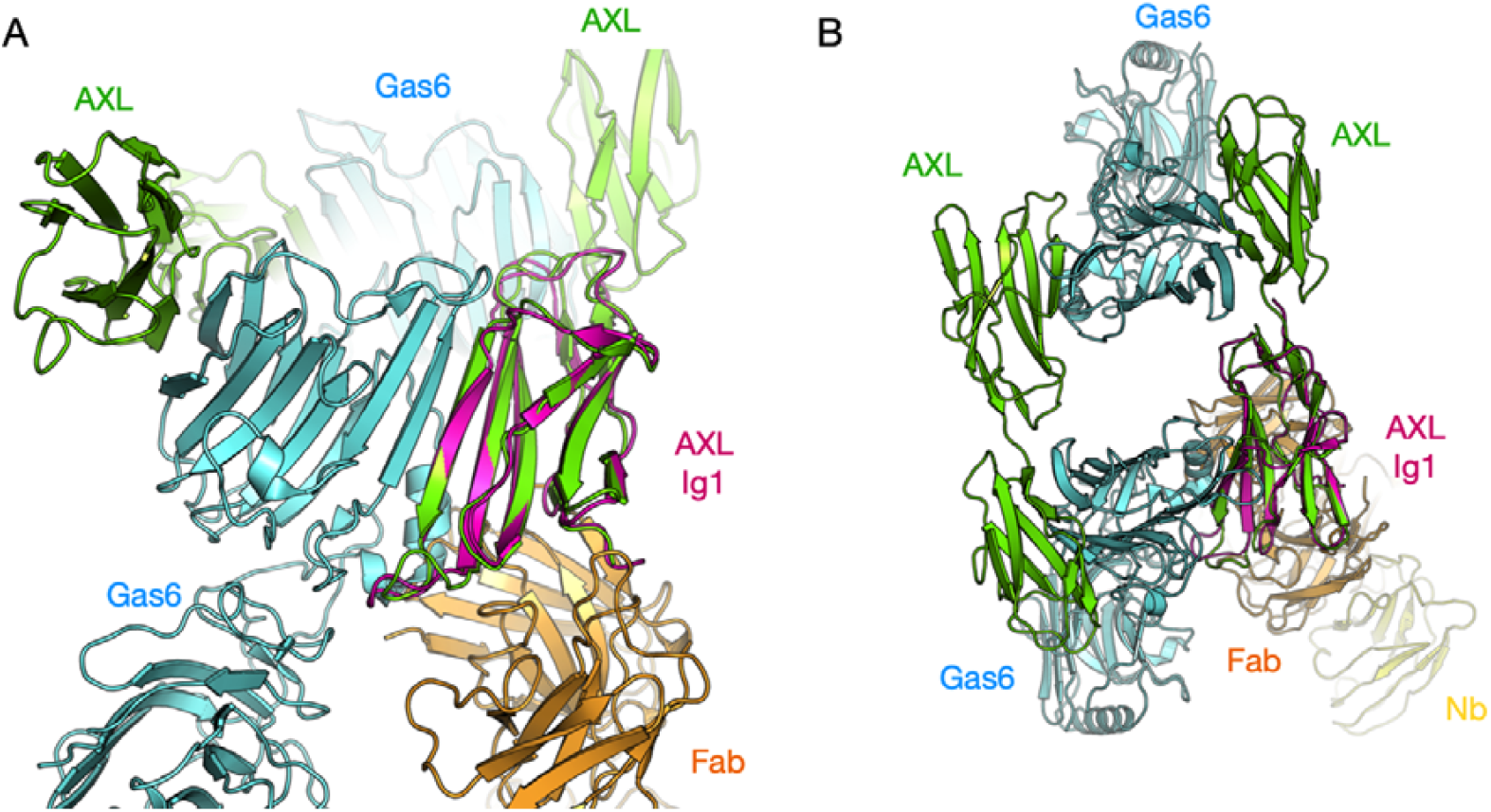
Comparison of the cryo-EM model (magenta/orange/yellow) to the GAS6-AXL complex crystal structure (green/cyan). **(A)** A view from the side of the AXL-GAS6 heterotetramer. The tilvestamab Fab binding overlaps slightly with the GAS6 binding site on AXL Ig1. **(B)** The top view of the heterotetramer indicates that tilvestamab binding does not directly interfere with oligomerisation, but its effects could be mediated through conformational changes.

## Conclusion

We have presented the first experimental results on the 3D structure of the complex between the AXL extracellular domain and tilvestamab. AXL alone had already earlier been shown to be highly extended and flexible, and in line with this, it remains elongated in the presence of tilvestamab and its Fab fragment, supporting the binding of tilvestamab to the first Ig domain of AXL. The structural data, which highlight the importance of experimental structural data especially for antigen-antibody complexes, will allow further development of tilvestamab and other specific AXL binders for therapeutic and diagnostic applications.

## Acknowledgements

This work was funded by an Industrial PhD grant from the Research Council of Norway (311399). We acknowledge the use of the Core Facility for Biophysics, Structural Biology, and Screening (BiSS) at the University of Bergen, which has received infrastructure funding from the Research Council of Norway through NORCRYST (grant number 245828) and NOR–OPENSCREEN (grant number 245922). This work has been supported by iNEXT-Discovery, grant number 871037, funded by the Horizon 2020 program of the European Commission. Access to iNano cryo-EM facilities at Aarhus University was thereby supported by iNext-Discovery project numbers 12416, 20077, and 25639. We thank Remy Loris (VIB-VUB Center for Structural Biology & Vrije Universiteit Brussel) for providing access to JEM-1400 electron microscope. We further acknowledge provision of synchrotron beamtime and excellent on-site support for SAXS experiments at Soleil synchrotron and MAX-IV.

