## Supplementary Figure 1 for "Structural analysis of tilvestamab in complex with AXL"

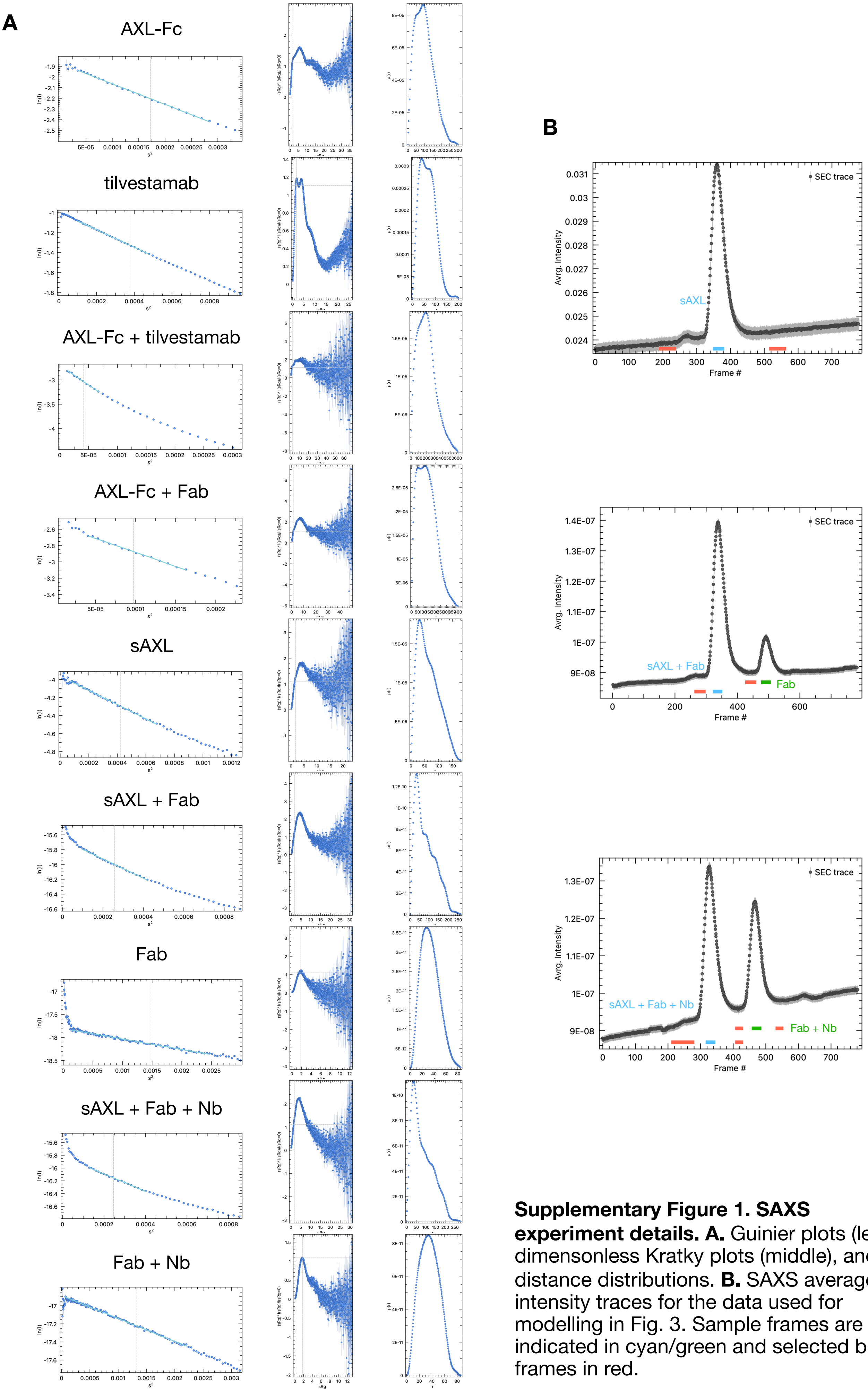


**Supplementary Figure 1. SAXS experiment details. A.** Guinier plots (left), dimensonless Kratky plots (middle), and distance distributions. **B.** SAXS average intensity traces for the data used for modelling in Fig. 3. Sample frames are indicated in cyan/green and selected buffer frames in red.
